# Complexes Formed by Mixing Tumor-derived Extracellular Vesicles with Polymeric Surfactants for Personalized Therapeutic Vaccine

**DOI:** 10.1101/2022.11.05.515294

**Authors:** Wenfeng Zeng, Hongjian Tian, Zihao Wang, Siqi Li, Lingtao Jin, Wei Liang

**Affiliations:** Protein and Peptide Pharmaceutical Laboratory, Institute of Biophysics, Chinese Academy of Sciences, Beijing, 100101, China; University of Chinese Academy of Sciences, Beijing, 100864, China; Department of Molecular Medicine, University of Texas Health Science Center at San Antonio, San Antonio, TX, 78229, USA

**Keywords:** Tumor-derived Extracellular Vesicles, Personalized Therapeutic Vaccine, Polymeric Surfactant, Immunosuppression

## Abstract

The personalized therapeutic vaccine is an ideal weapon to eliminate tumors. However, the core steps of manufacturing personalized cancer vaccines are identifying tumor-specific antigens (TSAs, also called neoantigens) and HLA epitope prediction, which is time-consuming and labor-intense. Tumor-derived extracellular vesicles (TEVs) are alternative sources of neoantigens. However, the immunosuppressive nature of TEVs limits their application in such immunotherapy. In this study, we present a new strategy to maintain neoantigens in TEVs and diminish the immunosuppression by deconstructing the structure of TEVs with polymeric surfactant polyethylene glycol-phosphatidylethanolamine (PEG-PE). Together with adjuvant MPLA, the newly formed micelle-like complexes compose a therapeutic vaccine (MLC-V). Results show that MLC-V is capable of eliciting neoantigen-specific T-cell responses, restoring TEV-induced immunosuppression, and preventing lung metastasis of murine melanoma. MLC-V also exhibits outstanding anti-tumor efficacy in multiple tumor models. MLC-V can be used as a personalized therapeutic vaccine in a mimetic pre-clinical MC38 model and the anti-tumor effect of MLC-V was synergistically enhanced by PD-1 mAb. Taken together, the present study demonstrates a time-saving, low-cost, and simplified strategy to produce personalized therapeutic vaccines based on MLC-V platform technology.

## Introduction

It has been convinced that somatic mutations in tumor cells can lead to tumor specific antigens (TSAs) generation, also called neoantigens, which can be recognized by the host immune system and assist adaptive immunity to kill tumor cells^[1,2]^. As vast majority of cancer mutations are unique to the individual patient, personalized approaches are needed. Numerous clinical trials have attempted to harness this mechanism of inducing anti-tumor response by using personalized neoantigen-based vaccines^[1,3–5]^. The synthesis strategy for personalized neoantigens mainly depends on the integration of next-generation sequencing and high-resolution and tandem mass spectrometry techniques. In brief, matched patient tumor tissue and adjacent normal tissue are subjected to whole-exome sequencing and RNA-Seq to identify expressed nonsynonymous somatic mutations. Then the adhesion affinity to HLA of these mutated neoepitopes is predicted by computer algorithms^[6,7]^ and expansion of mutant neoantigen-specific autologous T cells isolated from blood or tumor of the same patient. Ex vivo–expanded T cells are then infused back into the cancer patient^[4]^. Alternatively, the predicted neoantigens used to produce neoantigens-based vaccines. Despite the identification of personalized tumor neoantigens is practicable, it is still a multistep process which requires significant time, cost and labor to accomplish. The long-time process may cause patients enrolled miss the best therapeutic window due to the rapidly dynamics of neoantigens, which limits the application of personalized neoantigen-based immunotherapies in clinical practices^[8]^.

Extracellular vehicles (EVs) are membranous vesicles released by nearly all types of cells into extracellular space, including microvesicles, exosomes and apoptotic bodies, based on their sizes and origins. EVs carry an array of molecules (proteins, lipids, nucleic acids) that reflect the identity and activity of their cell-of-origin^[9,10]^. Tumor derived extracellular vesicles (TEVs) play a critical regulatory role between immune cells and tumor cells, and also TEVs carry both tumor-associated antigens (TAAs) and tumor specific antigens (TSAs, also known as neoantigens) inherited from parental tumor cells, such as HER-2/neu, EGFR2, CEA, MART-1, gp100, TRP-1, mesothelin, and members of the HSP family including HSP70 and HSP90^[11–15] [16]^. The tumor antigens in TEVs could be transferred to dendritic cells (DCs), and induce antigen-specific cytotoxic T lymphocyte (CTL) responses both *in vitro* and *in vivo*^[17]^. Additionally, TEVs elicited more effective protection against autologous tumors than irradiated tumor cells, apoptotic bodies, or tumor lysates *in vitro*^[18,19]^ suggesting TEVs are an ideal source of tumor antigens. More importantly, the isolation of TEVs is more convenient than a long-time and high-cost neoantigen production process, which includes WES, neoantigen identification, verification and preparation.

However, TEVs also show immunosuppressive capacity, resulting in either tumor progression or regression, which restricts the application of TDEs in therapeutic vaccines. TEVs induce T cell apoptosis via membrane-bound CD95 ligand (FasL) ^[20]^, and inhibit T cells function through TEV membrane-bound TGF-β^[21]^ and PD-L1^[22,23]^. Meanwhile, antigen presenting cells, like DCs, are paralyzed by TEVs via fatty acids-induced PPARα activation^[24]^ and PGE2-induced CD73 upregulation^[25]^. To overcome the immunosuppression of TEVs, modifications were reported to equip TEVs with immuno-stimulators, such as CD40 ligand enriched in exosomes heat-stressed 3LL Lewis lung tumor cells to promote DCs maturation^[26]^ and TAAs plus a TLR4 adjuvant to enhance tumoricidal T cells activity through the N-terminus domain of HMGN1 (N1ND)^[27]^. Although effective, such strategies are limited by the requirement to modify TEV producer cells, which often is time-consuming and challenging^[28]^. Of note, most of the above-mentioned strategies do not interrupt the structural integrity of TEVs, implying that the immunosuppressive molecules still exist within TECs, leading to poor outcomes.

Here we propose a novel strategy to utilize TEVs as a therapeutic vaccine via deconstructing the structure integrity by mixing TEVs with polyethylene glycol-phosphatidylethanolamine (PEG-PE) and MPLA (as adjuvant). The mixture represents a nano-scale complexes with micelle-like shape, tumor antigens remained and immunosuppression diminished, thus we name the new complexes as micelle-like complex vaccine (MLC-V). MLC-V shows robust anti-tumor effect in multiple murine tumor models, capability of eliciting antigen-specific CTL responses and inhibiting lung metastasis of melanoma cells. And most importantly, MLC-V strategy exhibited the potential to be used in personalized therapeutic vaccine preparation in a murine colon carcinoma model.

## Materials and Methods

### Mice

Female C57BL/6 mice (6–8 weeks old) were purchased from Vital River Laboratory Animal Technology Co. (Beijing, China). OVA257-264-specific TCR transgenic mice C57BL/6-Tg (TcraTcrb)1100Mjb/J (OT-I) were purchased from Jackson Laboratories (Bar Harbor, ME, USA). All the mice were housed under pathogen-free conditions in the animal care facilities at the Institute of Biophysics, Chinese Academy of Sciences. All animal experiments were approved by the Institutional Laboratory Animal Care and Use Committee at the Institute of Biophysics, Chinese Academy of Sciences.

### Cell lines and Reagents

Murine breast cancer 4T1, cervical carcinoma TC-1, colon carcinoma MC38 and melanoma B16/F10 were cultured in 5% CO2 and maintained in RPMI 1640 or DMEM medium supplemented with 10% FBS (BI, Isreal), 100 U/ml penicillin, and 100 μg/ml streptomycin. 4T1, TC-1 and B16/F10 were obtained from ATCC, MC38 were obtained from the laboratory of Yangxin Fu, and the test for mycoplasma infection were negative. Anti-mouse PD-1 (RMP1-14) antibody was purchased from BioXcell.

### Isolation of TEVs

Tumor cells were cultured in DMEM or RPMI 1640 media supplemented with 10% TEVs - depleted FBS in which bovine TEVs were removed by ultra-centrifugation at 100,000 g for 2 hours. Supernatant of tumor cells culture was collected 48 hours after cell reached 80% confluence. Then the supernatant was centrifuged at 4000 rpm for 2 hours to remove cell debris, followed by 4000 rpm centrifuge for 30 min using 100 KDa MWCO filter to collect the TEVs - concentrated solution. The TEV quick extraction solution (EXOTC10A-1, SBI) was added to the TEVs - concentrated solution at 1:5 volume ratio and stored at 4°C for at least 12 hours. The precipitation was dissolved with PBS and stored at −80°C. The protein content of TEVs was determined by BCA protein assay kit (ThermoFisher).

### Preparation of MLC-V

The MLC-V was prepared by film-rehydration method ^[55]^. Briefly, PEG-PE (Lipoid, Newark, NJ, USA) dissolved in 20 mg/ml with chloroform (Sigma, St Louis, MO, USA). MPLA (Avanti Polar lipids, Alabaster, AL, USA) was dissolved in chloroform/methanol (2:1).to 5 mg/ml. PEG-PE solution and MPLA were mixed in a mass ratio of 100:1, then the mixture of PEG-PE: MPLA: TEVs is MLC-V.

### Cryo-Transmission electron microscopy (Cryo-TEM) and Dynamic Light Scattering

PEG-PE (Lipoid, Newark, NJ, USA) micelle is prepared as described before^[55]^. For cryo-TEM, the protein concentration of TEVs, PEG-PE or PEG-PE mixed with TEVs (PP/EXO) samples were all set on 5 mg/ml. And the samples were transferred to a glow-discharged R1.2/1.3 Quantifoil Holey carbon film grid and incubated for 30 s. Grids were blotted for 6s at level 4 force and immediately plunge frozen into nitrogen-cooled liquid ethane using a Vitrobot Mk IV (Thermo Fisher Scientific). The carbon mesh was observed under a frozen transmission electron microscope (Talos L120C at 120kV).

For size distribution and particle dispersal index (PDI), TEVs, PEG-PE and PP/TEVs were determined by DLS using a Malvern Zeta-sizer Nano ZS (λ = 633 nm, Zetasizer Nano ZS, Malvern, UK) at 298 K. The micelles were diluted to 1 mg/ml with saline. The samples were diluted to 0.25 mg/ml of TEVs with PBS.

### Transfection and stable cell line construction

293T cells were transfected with 7 μg of the mixed plasmids (phage-EF1α-PD-L1-IRES: psPAX2: pMD2.G = 4:3:1), and with 28 μl Lipofectamine 3000 (Invitrogen) in 1 ml Opti-MEM mediem for the production of lentivirus. Then, MC38 cells were transduced with the lentiviral vector to overexpress PD-L1 gene. Transfection efficiency was analyzed by flow cytometry and PD-L1 expression of TEVs was detected by Western Blot.

### LC-MS and data analysis

MC38 cells and their secreted EVs (triplicate each) were lysed on ice for 2 h in lysis buffer (7 M urea, 2% SDS, and 1 × protease inhibitors). The lysis supernatant was quantified by BCA Protein Assay Kit (ThermoFisher). Protein precipitation was performed by adding Dithiothreitol (DTT) at 37 °C for 1 h, then IAA was added in a volume of fivefold than DTT for another 40 minutes. After incubation, precipitated material was pelleted by centrifuging for 1 h at 15,000 g at 4 °C. The pellet was washed with acetone and let dry at room temperature. trypsin for lysis. Next, the pellet was lysed with typsin. All the samples were mixed and then incubated overnight at 37 °C. After the incubation, trypsin activity was quenched by adding 10% formic acid. Before analysis by LC-MS/MS, each sample was desalted using Pierce C18 spin columns, after which each sample was dried and reconstituted in 50 mL water with 0.1% formic acid and transferred to a high-performance liquid chromatography (HPLC, EASY-nLC 1000, Thermo Scientific, USA) vial for analysis.

The tryptic peptides were loaded onto an Acclaim PepMap 100 C18 trapping column(C18, 1.9 μm, 75 μm*20 cm; Thermo Fisher) at a flow rate of 200 nL/min. MS/MS spectra were measured by Orbitrap Fusion Lumos (Thermo Scientific, USA) across an m/z range of 350–1,600 Da at a resolution of 60,000, and fragmented in the HCD cell using a collision energy of 30. Proteins were first identified and quantified (label-free) using Proteome Discoverer2.4 (Thermo Fisher) and the Sequest HT algorithm combined with the Target Decoy PSM Validator.

### Western Blot Analysis

Samples were lysed in RIPA buffer (Beyotime) with the addition of proteasome inhibitors (Sigma-Aldrich). Western Blot was performed as described in the commercially available antibody protocols.

### T cell proliferation assays

CD8^+^ T cells were harvested from OT-I transgenic mice and purified by magnetic beads according to the manufacturer’s instructions (Introvigen). 5×10^4^ BMDCs were pulsed with 1 mg/ml endotoxin-free chicken egg ovalbumin (OVA, Hyglos GmbH, Germany) in the presence of 50 μg/ml MC38 TEVs or MLC-V. After 16 hours of incubation at 37°C, BMDCs were washed. Purified OT-I CD8^+^ T cells were labelled with CFSE, according to the manufacturer’s instructions (Invitrogen), and 4×10^5^ T cells were incubated with the BMDCs (Ratio of DC: T = 1:8). After 3 days co-incubation, proliferation of CD8^+^ T cells were analyzed by FACS.

### IFN-γ ELLISpot assay

IFN-γ ELLISpot assays were performed using the Mouse IFN-γ ELISPOT Set Kit (BD Biosciences)) according to manufacturer’s instructions. Plates were coated overnight at 4 °C with anti-IFN-γ Capture Antibody diluted in Coating Buffer, washed with Coating Buffer and blocked with Cell culture medium (RPMI 1640) containing 10% Fetal Bovine Serum (FBS) and 1% Penicillin-Streptomycin-L-Glutamine containing 1% penicillin/streptomycin (GIBCO) for 2 h at room temperature.

For *ex vivo* ELISpots, lymphocytes from inguinal lymph nodes were plated with 3×10^5^ cells per well. Peptides were added to ELISpot wells at 20 μg/ml per peptide. Each plate included a healthy donor positive and negative control with the Phorbol myristate acetate (PMA, InvivoGen) to confirm reagent performance. Plates were incubated 48 h at 37°C. Plates were washed 3 times using PBS with 0.05% Tween 20 and detection antibody diluted in Dilution Buffer was added to wells for 2 h. After washing 3 times with PBS with 0.05% Tween 20, Avidin-HRP was diluted in Dilution Buffer and added to wells for 30 min. Plates were washed with both PBS with Tween 20 and PBS 3 times. AEC substrate-chromogen (BD Biosciences) was then added for 5-20 min. Plates were rinsed with deionized water 3 times and allowed to dry at room temperature overnight. Spots were imaged and enumerated using an Immunospot analyzer (CTL analyzer LLC).

### Murine Melanoma Lung Metastases Model

50 μg B16F10 TEVs or MLC-V (500:5:50 μg) based on B16F10 or PD-L1^OE^B16F10 EVs were injected intravenously to the mice for 3 doses within a week. Then lung metastases were established by intravenous injection of 5×10^4^ B16F10 cells into mice. 3 weeks later, the mice were sacrificed and the metastatic nodules on lungs were counted.

### Therapeutic Applications of MLC-V in Multiple Murine Tumor Models

For the therapeutic tumor vaccination studies, 6–8-week-old female C57BL/6 mice were subcutaneously inoculated at the right flank with 2.5×10^5^ cells per mouse (in MC38 tumor model), or with 5×10^4^ cells per mouse (in TC-1 tumor model), or with 2.5×10^5^ cells per mouse (in B16-F10 tumor model). After the tumor was established, the mice were vaccinated with 50 μg/ml TEVs and MLC-V based on the TEVs for 3 doses at the interval of one week. The tumor volume in this study was calculated by the equation: tumor volume = length × width^2^ × 0.5. Animals were euthanized and seen as dead when the tumor volume reached 1000 mm^3^.

For personalized therapeutic vaccine strategy, the C57BL/6 mice were subcutaneously inoculated at the right flank with 2.5×10^5^ MC38 cells per mouse, and the tumor tissues were dissected on day 9. We isolated and purified tumor cells by Tumor Cell Isolation Kit (Miltenyi Biotec) and cultured until 4^th^ generation. The tumor cells were then subcutaneously transplanted on C57BL/6J mice (wild type 6–8-week-old female) to establish a new tumor model. Meanwhile, the supernatant of 4^th^ generation tumor cells was collected to isolate Tumor EVs. On day 4 after the transplantation, MLC-V(Tumor) based on Tumor EVs was vaccinated to the mice bearing tumor for continuously 2 weeks, once a week. And the MLC-V based on primary MC38 cell line derived EVs was injected as control.

For anti-PD-1 combination experiment, vaccinated animals were intraperitoneally administrated with 100 μg anti-PD-1(RMP1-14, BioXcell) per mouse or PBS at days 11 and 14 post tumor injection.

### Statistical analysis

Statistical analysis was performed using GraphPad Prism statistical software, Version 8 (GraphPad Software Inc, San Diego, CA, USA). Differences between groups were calculated using Student t test. Log-rank test was performed for survival analysis. A value of P < 0.05 was considered statistically significant (*P < 0.05; **P < 0.01; and ***P < 0.001; **** < 0.0001)

## Results

### PEG-PE Deconstructs TEVs to Form Micelle Like Complexes

TEVs contain TSAs^[29]^, and to verify these findings in our experimental settings, we compared the proteins in murine colon carcinoma cell line MC38 with those in MC38-secreted EVs by MS/LS. There were 6218 proteins identified in parent MC38 cells and 570 proteins in MC38 EVs. Among the 570 proteins in TEVs, 432 proteins overlapped with cellular proteins, accounting for nearly 76% of proteins of MC38 EVs. Also, three identified neoantigens, Adpgk^[6]^> Rpl18^[30]^ and Cspg4^[31]^ were detected in MC38 EVs (Fig.1A and B).

**Fig.1.**
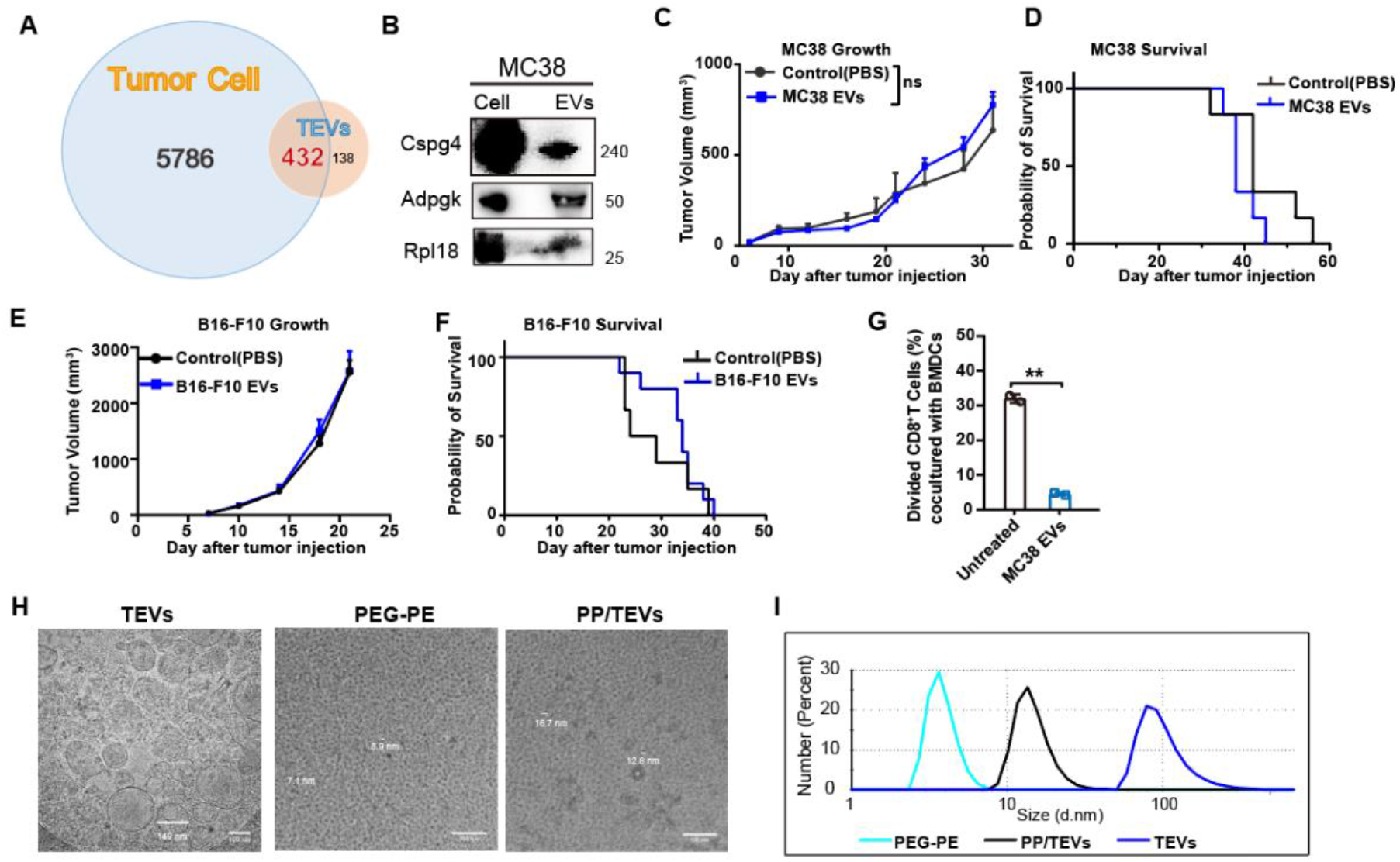
PEG-PE Deconstructs TEVs to Form Micelle Like Complexes. **A**, Venn diagram of proteins identified by proteomics from MC38 cells and MC38-derived EVs. **B**, MC38 TSAs Rpl18, Adpgk, Cspg4 were detected by western blot. **C-F**, Anti-tumor effect of MC38 EVs and B16-F10 EVs on tumor-bearing mice. C57BL/6 mice were inoculated subcutaneously with 2.5×10^5^ MC38 cells or 2.5×10^5^ B16F10 cells, then immunized with MC38 EVs (50 μg/mice) or B16-F10 EVs (50 μg/mice) on days 4, 11 and 28. Tumor growth (**C**) and survival curves (**D**) of MC38 model, tumor growth (**E**) and survival curves (**F**) of B16-F10 model were shown. Data are shown as the Mean ±SEM from a representative experiment of 2–3 independent experiments with n = 6. **G**, The effect of MC38 EVs on DCs function of priming CD8^+^ T cells. Data are shown as the Mean ±SEM (n=3). ns, no significance. **, p<0.01 by Student t test. **H**, **I**, Cryo-electron microscopy (**H**) and dynamic light scattering analyses (**I**) of PEG-PE micelle, TEVs, and PEG-PE mixing with TEVs. Scale bar, 100 nm.

However, TEVs neither reduced tumor growth nor extended the life span of tumor bearing mice in colon carcinoma (MC38) or melanoma (B16-F10) model (Fig.1 C-F), indicating that it is not a practicable to use TEVs without modifications as a therapeutic vaccine. The immunosuppression of TEVs may underscore these results, since the *ex vivo* study showed that TEVs inhibited the priming effect of bone marrow dendritic cells (BMDCs) on T cells (Fig.1 G).

To overcome the immunosuppressive nature of TEVs, we propose a new strategy to deconstruct TEVs structure by PEG-PE, a polymeric surfactant. The cryo-TEM images demonstrated that the mixture of PEG-PE and TEVs resembled PEG-PE micelles, but not the typical oven-like structures of TEVs (Fig.1 H), suggesting structure integrity of TEVs was lost and a new micelle-like complexes were reformed with a uniform particle size (~20 nm) distribution. And, this observation was further supported by DLS analysis, that the average size of PEG-PE treated TEVs were much smaller than TEVs, but comparable to PEG-PE micelles (Fig.1 I). Thereupon, we added MPLA as adjuvant to the system to produce a new micelle-like complexes vaccine (MLC-V).

### MLC-V Elicits Neoantigen-Specific Cytotoxic T Lymphocyte Responses

To explore whether MLC-V is capable to transfer antigen to DC cells and elicit neoantigen specific cytotoxic T lymphocyte responses in mice, we prepared MLC-V using MC38 EVs. First, we established an OVA, the paradigm antigen, expressing MC38 cell (MC38-OVA) line and collected the EVs. Western blot results showed that OVA was presented in MC38-OVA EVs (Fig.2 B). Then, T-Select H-2K^b^ OVA Tetramers were employed to detect antigen (OVA) specific T cells from lymph nodes of mice immunized with MC38-OVA EVs or MLC-V based on MC38-OVA EVs (Fig.2 A). Compared with TEVs, OVA Tetramers positive CD8^+^ T cells percentage is obviously higher in MLC-V immunized mice than TEVs group (Fig.2 B). Further, we vaccinated wild type C57BL/6J mice with MC38-EVs or MLC-V based on MC38 EVs. 7 days later, the lymphocytes were isolated and restimulated with MC38-specific neopeptides Rpl18 and Adpgk^[32]^, respectively. ELLISpot assay showed that, compared with MC38-EVs, the number of neo-peptides specific IFN-γ secreting cytotoxic T cells was about tenfold greater after MLC-V treatment (Fig.2 C), indicating MLC-V retains neoantigens from MC38 cells, and these neoantigens were successfully presented to T lymphocyte *in vivo*.

**Fig.2.**
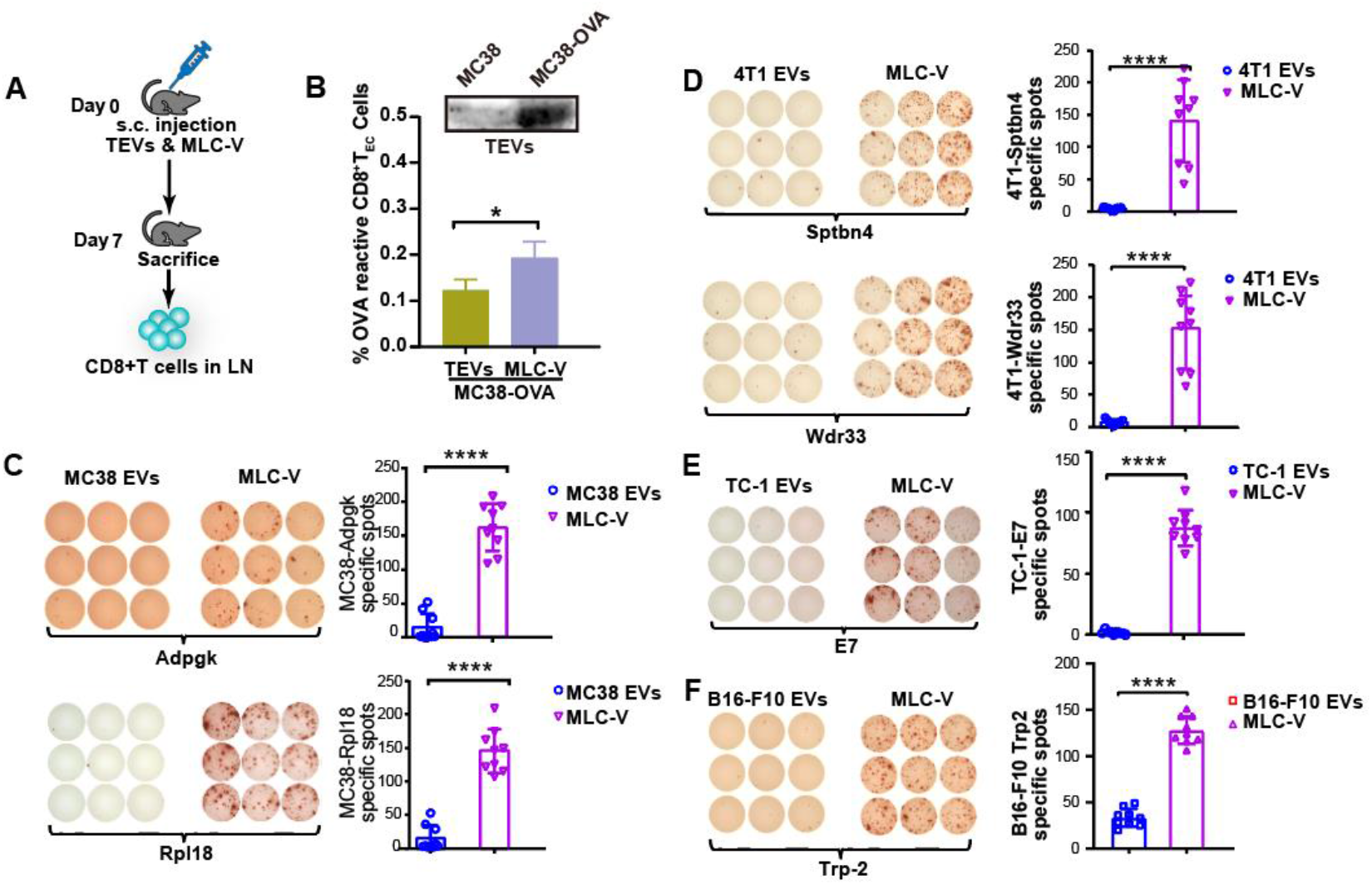
MLC-V Elicits Neoantigen-Specific Cytotoxic T Lymphocyte Responses. **A**, Schematic plot describing how OVA specific CD8^+^ T cells were detected *in vivo*. **B**, OVA was detected in MC38-OVA cell derived EVs by western blot (upper). Frequency of OVA tetramer specific CD8^+^ T cells in inguinal lymph nodes (lower, data were shown in Mean ±SEM; n = 3). **C-F**, **C**, ELLISpot analysis of Rpl18 (upper) and Adpgk (lower) specific CD8^+^T cells in inguinal lymph nodes of mice vaccinated with MC38 EVs and MLC-V based on MC38 EVs. **D**, ELLISpot analysis of Sptbn4 (upper) and Wdr33 (lower) specific CD8^+^T cells in inguinal lymph nodes of mice vaccinated with 4T1 EVs and MLC-V based on 4T1 EVs. ELLISpot analysis of E7 (E) and Trp-2 (F) specific CD8^+^T cells in inguinal lymph nodes of mice vaccinated with TC-1 or B16F10 EVs and MLC-V based on TC-1 or B16F10 EVs. Data are presented as IFN-Y spot-forming units (SFUs) per 3×10^5^ splenocytes (Mean ±SEM; n = 9) from a representative experiment of 2–3 independent experiments *, P<0.05; ****, P<0.0001 by unpaired Student’s 2-tailed t test.

Next, we asked whether the effect of MLC-V on eliciting neoantigen-specific CTL responses was universal. To address this issue, we used the same strategy to evaluate EVs from murine triple-negative breast cancer 4T1, HPV-related cervical cancer TC-1 and Melanoma B16-F10. Lymphocytes were restimulated with 4T1-specific neo-peptides Sptbn4 or Wdr33 (Fig.2 D), TC-1-specific neo-peptide E7 (Fig.2 E), and BF16-F10-specific neo-peptide Trp-2 (Fig.2 F). Results demonstrated that all tested MLC-Vs displayed dominant responses toward their neo-epitopes (Fig.2 D-F).

### MLC-V Reverses TEV-Induced Immunosuppression

It is well known that DCs orchestrate anti-tumor immunity and effector T cells are the main executors in killing tumor cells. Thus, in order to investigate whether TEV-induced immune dysfunction could be overcome by MLC-V, *in vitro* DC priming T cell assay and T cell proliferation assay were both employed. Our previous study showed that the ability of TEV-treated BMDCs to prime OVA antigen specific T cell activation was strongly undermined ^[33]^. However, in the present research, compared with TEV treatment, MLC-V (based on MC38 EVs) treated BMDCs restored the capability to prime OT-I CD8^+^ T cells (Fig.3 A).

**Fig.3.**
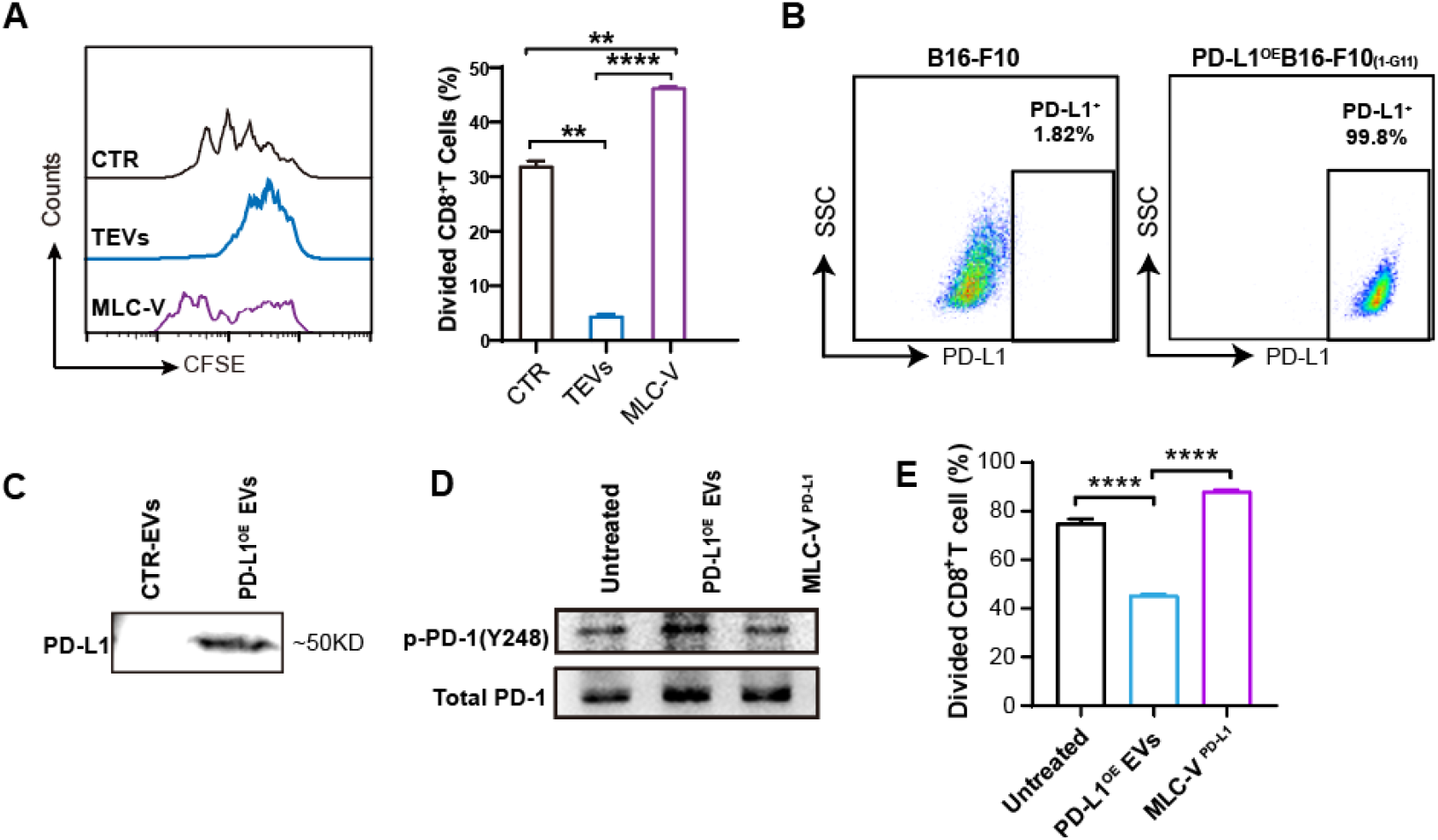
PEG-PE Reverses TEV-Induced Immunosuppression. **A**. The effect of MC38 EVs or MLC-V based on MC38 EVs on DCs’ function in priming OT-I CD8^+^ T cells. **B**. PD-L1 expression on B16F10 and PD-L1^OE^B16F10 cells by flow cytometry. **C**. PD-L1 expression on EVs of B16-F10 cells (CTR-EVs) and PD-L1^OE^B16F10 cells (PD-L1 ^OE^ EVs) by Western blot. **D**. Phosphorylated PD-1-Y248 and total expression of PD-1 in EVs treated T cells were detected by Western blot. **E**. The proliferation of T cells under different EVs treatment. Data are presented as Mean ±SEM (n = 3) from a representative experiment of 2–3 independent experiments. **, P<0.01; ****, P<0.0001 by unpaired Student’s 2-tailed t test.

Upregulating the expression of programmed death-ligand 1 (PD-L1) is one of the main strategies of tumor cells to evade immune surveillance^[34]^. Chen, et al reported that PD-L1 was evidently upregulated in exosomes from metastatic melanoma cells compared to those from primary melanoma cells, and enriched PD-L1 was also found in EVs from murine metastatic melanoma B16F10 cells^[35]^. Thus, we established a PD-L1 overexpressed B16F10 cell line (PD-L1^OE^B16F10) with lentivirus infection method (Fig.3 B). And PD-L1 was detected in purified PD-L1^OE^B16F10 EVs (PD-L1^OE^ EVs) (Fig.3 C). Then MLCV based on PD-L1^OE^ EVs (MLC-V ^PD-L1^) was prepared.

Moreover, once PD-1 on T cells ligated with exogenous PD-L1 either on the membranes of cells or EVs, the Y248 in cytoplasmic tail will be phosphorylated, which is indispensable for PD-1 mediated inhibitory function in T cells^[36]^. Higher PD-1-Y248 phosphorylation level was found in T cells treated with PD-L1^OE^ EVs compared to control EVs and MLC-V^PD-L1^ (Fig.3 D), indicating that PD-1 inhibitory signaling was blocked by MLC-V^PD-L1^. Next, we observed that T-cell proliferation is inhibited by the engagement of PD-L1^OE^ EVs, while the proliferation of MLC-V^PD-L1^ treated T cells was not affected (Fig.3 E). These results suggested PD-L1^OE^ EVs initiated strong PD-1 mediated suppression in T cells proliferation, and the interaction between PD-L1 presented on EVs and T cells was interrupted by MLC-V. And more importantly, the effect of MLC-V on both DCs and T cells further substantiate the propose that PEG-PE was capable to deconstruct the EVs’ structure.

### MLC-V Prevents Lung Metastasis of Melanoma

It has been reported that TEVs promoted metastasis of primary cancer cells by forming immunosuppressive niches in distal organs^[37–39]^, thus it would be interesting to ask whether MLC-V could prevent TEV-mediated distal metastasis. Intravenously (i.v.) injection of B16F10 cells would establish a passive lung metastasis model in wildtype C57BL/6 mice and higher level of PD-L1 on TEVs would further promote the metastasis. One week before B16-F10 injection, the mice were pretreated with either B16F10 EVs (CTR EVs), PD-L1^OE^ EVs or MLC-V ^PD-L1^ respectively. Results showed that PD-L1^OE^ EVs pre-treatment led to more metastasis than vehicle control and CTR-EVs, suggesting that TEVs, especially PD-L1 enriched TEVs were more capable to establish an immunosuppressive microenvironment for tumor cells metastasis. But MLC-V ^PD-L1^ pre-treatment reduced nearly 80% of lung metastasis compared with PD-L1^OE^ EVs treatments (Fig.4 A and B).

**Fig.4.**
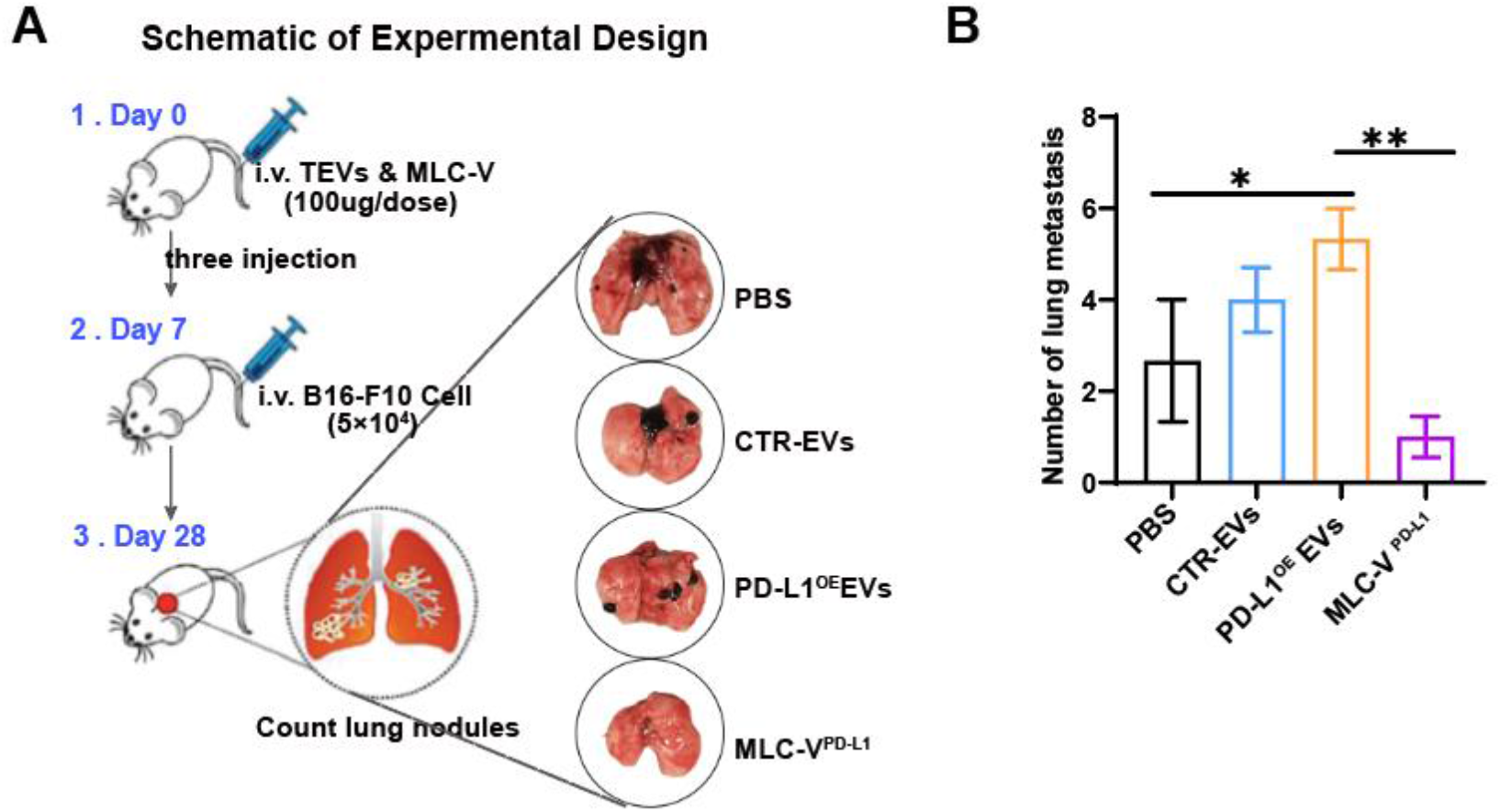
MLC-V Prevents Lung Metastasis of Melanoma. **A**. Schematic plot of lung metastasis model. Day 0, wildtype C57BL/6 mice were i.v. administrated with 3 doses of CTR-TEVs, PD-L1^OE^ EVs or MLC-V^PD-L1^ within one week, and 5×10^4^ B16-F10 cells were injected i.v. on day 7. Mice were sacrificed on day 28. Representative pictures of the lungs were showed in right panel. **B**. Number of lung metastatic nodules in each group. The data are shown as the Mean ±SEM (n ≥ 3 and were analyzed by one-way ANOVA. **, P<0.01; ****, P<0.0001.

### The Therapeutic Effect of MLC-V in Multiple Murine Tumor Models

Since MLV-C can induce CD8^+^ T cells dominant response toward tumor neo-epitopes, we aimed to therapeutically vaccinate mice with MLV-C after tumor inoculation. First, in murine colon carcinoma MC38 model, vehicle and MC38-EVs treatment did not affect tumor growth, while MLV-C treatment dramatically slowed the tumor growth (Fig.5 A) and extended lifespan (Fig.5 B), and resulting in one tumor free mouse. Similarly, in murine melanoma model (B16-F10) (Fig.5 C and D) and HPV-related cervical cancer model (TC-1) (Fig.5 E and FD), MLC-V also exhibited robust therapeutic effects.

**Fig.5.**
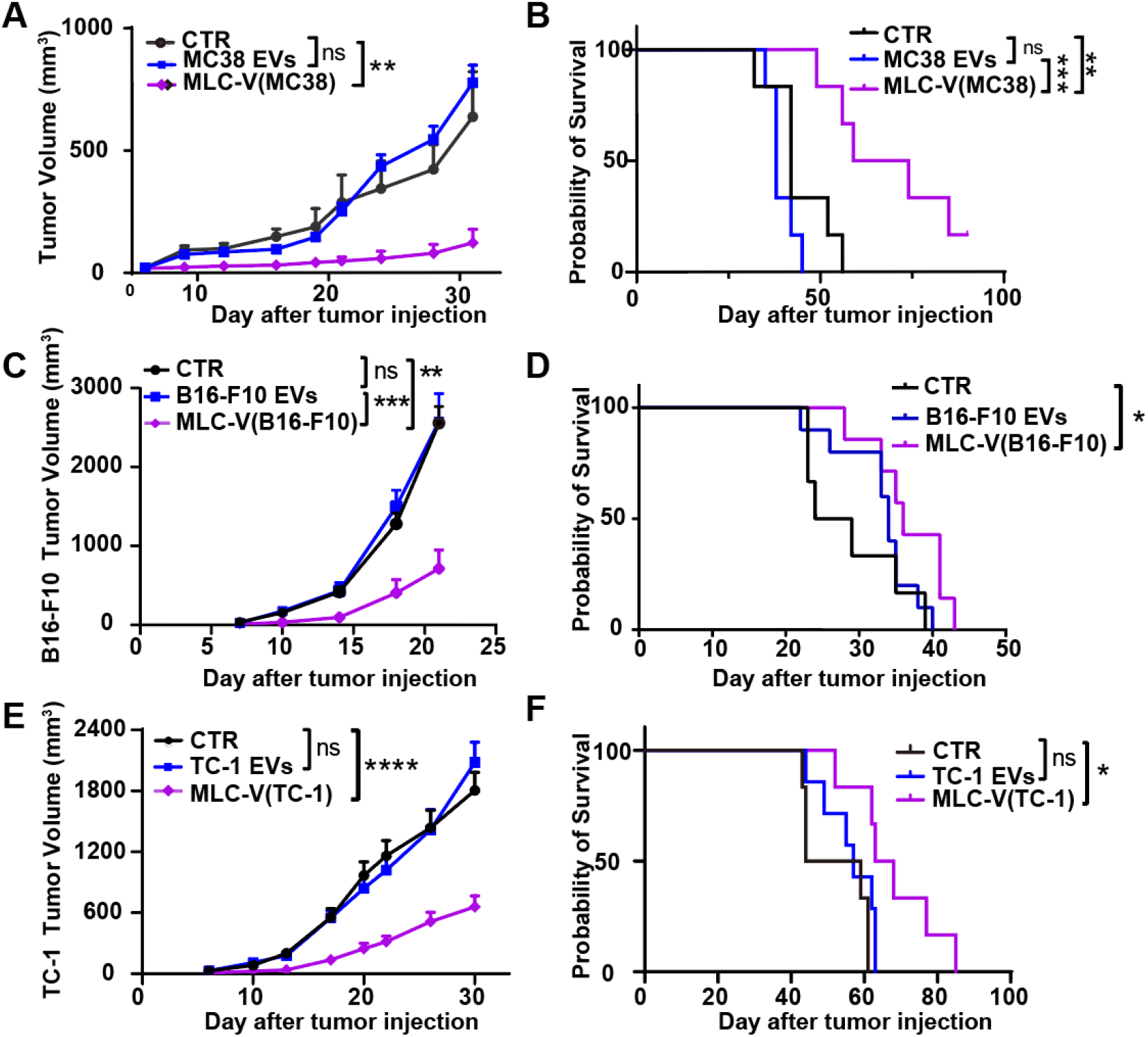
The Therapeutic Effect of MLC-V in Multiple Murine Tumor Models. **A**. The tumor growth and **B**. survival curves of MC38 model. **C**. The tumor growth and **D**. survival curves of B16F10 model. **E**. The tumor growth and **F**. survival curves of TC-1 model. Data are shown as the Mean ±SEM (n ? 5) from a representative experiment of 2–3 independent experiments. Tumor growth curves were analyzed by one-way ANOVA and survival curves were analyzed by Logrank test. ns, no significance; *, P<0.05; **, P<0.01; ***, P<0.001; ****, P<0.0001.

### Personalized Therapeutic Vaccine Simulation with MLC-V

In clinic, cancer cells are under immunoediting, which may lead to the alteration of neo-antigen spectrum. Does TEVs reflect these changes? If so, we reasoned that whether the MLC-V based on TEVs from those immunoedited cancer cells still capable of eliminating cancer cells? Since MC38 cell line has been proved to be a model for hypermutated/MSI colorectal cancer based on the landscape of genomic and transcriptomic characterization^[40]^. Thus, in immune competent mice, transplanted MC38 tumor will undergo genome mutation under the immunoediting pressure, indicating MC38 is a suitable model to mimic clinical cancer progress in patients.

Next, we designed a work flow to simulate biopsy of cancer patients and to purify TEVs from cancer tissues for MLC-V preparation (Fig.6 A). In brief, 7 days after primary MC38 cells were transplanted in the flank of mice, the tumor cells were isolated from dissected tumor tissue and cultured for 3 passages *ex vivo*. At the 4^th^ generation, the TEVs were isolated and purified from the supernatant (hereafter referred as Tumor EVs), and based on which MLC-V was prepared (MLC-V(tumor)). Meanwhile, the cells of 4^th^ generation were used to establish a new transplantable tumor model. 4- and 11-days post transplantation, the mice received 2 doses of MLC-V based on either purified tumor EVs or primary MC38 EVs. Interestingly, MLC-V(tumor) dramatically retarded tumor growth, and MLC-V(MC38) showed litter effect compared to the vehicle treated control group (Fig.6 B). This result strongly supported our hypothesis since the EVs from resected tumor cells must contain neoantigens from its parent cells and these neoantigens were different from those in EVs from primary MC38 cells. From another aspect, compared with traditional personalized vaccine production process, the preparation of MLC-V based personalized vaccine can shorten the time needed from 6-18 weeks^[41]^ to 2 weeks.

**Fig.6.**
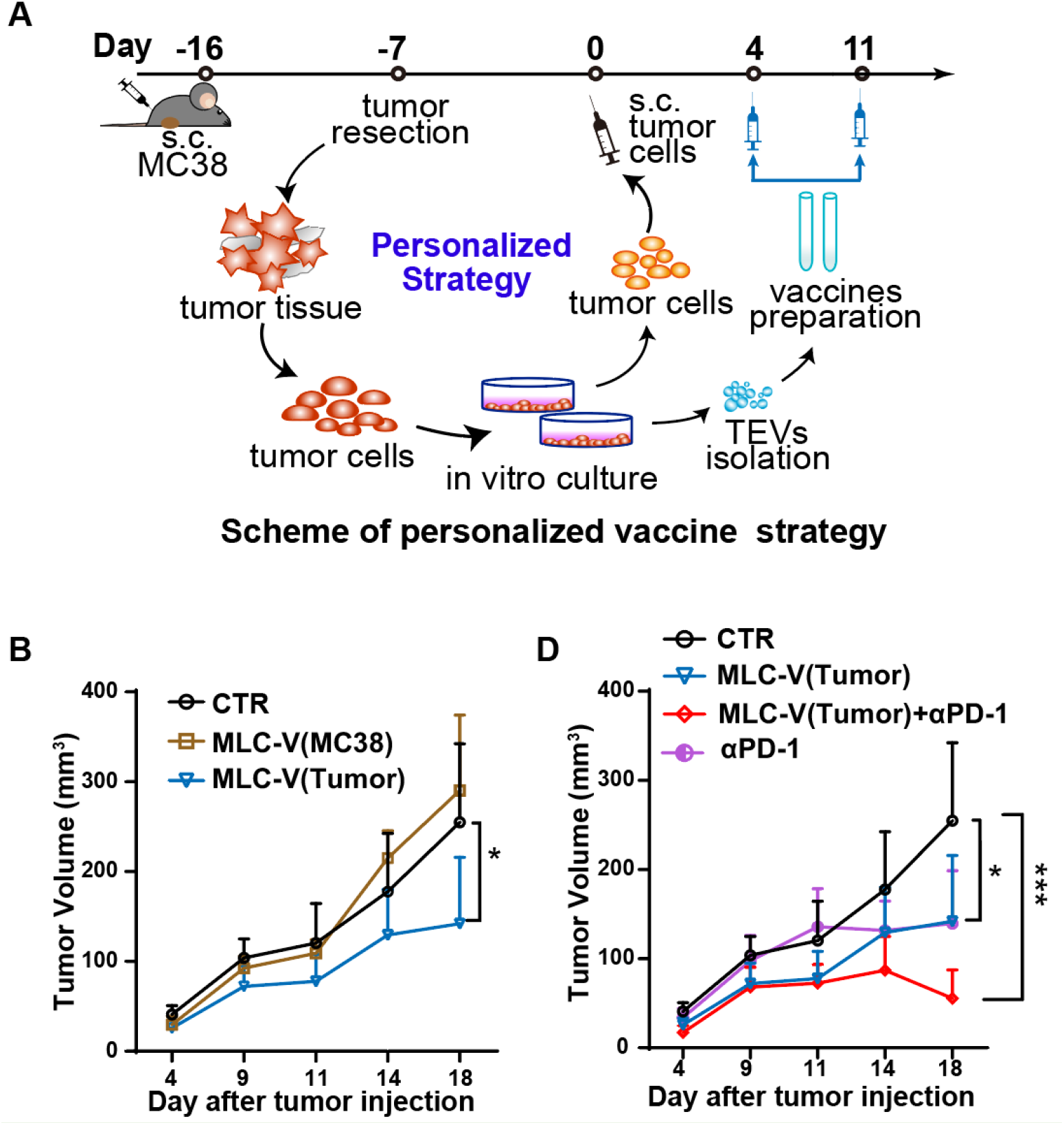
Personalized Therapeutic Vaccine Simulation with MLC-V. **A**. Schematic plot of personalized vaccine strategy simulates clinical scene. C57BL/6 mice were inoculated with MC38 cells on day −16, and 9 days later the tumor tissues were resected and the tumor cells were purified until 4th generation. The new tumor EVs were isolated from the supernatant of 4^th^ generation tumor cells. Meanwhile, the newly-obtained tumor cells were transplanted on C57BL/6J mice to build a new tumor model. In this new tumor model, MLC-V(Tumor) based on tumor EVs and MLC-V(MC38) were vaccinated to the tumor bearing mice on day 4 and 11. And anti-PD1 mAb was i.p. administrated on day 11 and 14. **B**. The tumor growth of vehicle (CTR), MLC-V(MC38) and MLC-V(Tumor) treated groups. **C**. The tumor growth curve of vehicle (CTR), MLC-V(Tumor), MLC-V(Tumor) combined with anti-PD1 and anti-PD-1 treated groups. The data are shown as the Mean ±SEM (n ≥ 6) and were analyzed by one-way ANOVA. ns, no significance; *, P<0.05; ***, P<0.001.

Additionally, therapeutic vaccines usually need to combine other therapies to gain ideal prognosis. In clinic, checkpoint inhibitors, especially monoclonal antibodies against PD-1/PD-L1 have become front line choices in multiple cancers treatment, including colon carcinoma^[42–44]^. Therefore, we investigated whether PD-1 mAb could synergize the therapeutic efficacy of MLC-V in the above-established model. Results showed that the combination of MLC-V (Tumor) and PD-1 mAb significantly enhanced antitumor efficacy as compared to the monotherapy alone (Fig.6 C). The present study indicates the potential of MLC-V strategy to facilitate next generation of personalized therapeutic vaccine and the compatibility of MLC-V to other anti-tumor therapeutics.

## Conclusion

Although recent advances in next-generation sequencing and computational predictions have been successfully used to identify neoantigens and to make individualized therapies feasible in clinical practices^[45]^. The biggest obstacle, however, is the rapid and accurate identification of immunogenic neoantigens. The procedure involves identification of potent immunogenic neoantigens by whole exome sequencing (WES) on matched tumor cells and adjacent normal cells^[46]^, followed by validation and analysis via RNA sequencing^[47]^, complicated HLA-allele typing and match^[48]^, predictions of tumor antigenic epitope-HLA-allele interactions and identification of cancer-specific MHC I-associated peptides (MAPs)^[49]^. Finally, the immunogenicity of synthesized neoantigen peptides must be verified by autologous APCs (antigen-presenting cells) stimulating T cell assay. Therefore, it is time-consuming and labor-intense procedure hindering prompt therapeutic cancer vaccines application^[50]^. During the long process, more than 6 weeks as reported^[2,4,41]^, tumor may progress rapidly and under the stress of immunoediting or chemotherapy, radiotherapy the spectrum of neoantigen shifts. In addition, current successful rate of neoepitope predictions remains poor, and an effective personalized vaccine needs up to 10-20 predicted neoantigens to be targeted ^[3]^, resulting in a complicated and high-quality neoantigens production.

Recently, TVEs-based immunotherapy has emerged as a new track in cancer immunotherapy owing to the nature that TEVs carry antigens direct from the parent tumor cells, and the capability to induce neoantigen-specific cytotoxic T lymphocyte (CTL) responses. However, TEVs also harbor immunosuppressive nature, resulting in tumor progression or regression^[39,51]^, which restricts the application of TEVs in therapeutic vaccines. Here, we use TEVs as a source of tumor neoantigens, and propose a new strategy of deconstructing the structure of TEVs by polymeric surfactants PEG-PE. Our results validated that PEG-PE transformed TEVs into micelle like complexes with the uniform particle size (~20 nm), and the addition of adjuvant MPLA made a new micelle-like complexes vaccine (MLC-V). MLC-V overcomes TEV-induced immunosuppression, not only reverses overall dysfunction to DCs, but also restores TEV-bound PD-L1 induced T cell suppression. Other TEV related molecules also induce immunosuppression, such as miRNAs, lipids or other membrane-bound proteins (CD73, FasL, etc), which was not verified by the present study that whether MLC-V could also overcome these molecules induced immunosuppression.

Then, we mimicked the clinical settings to harvest TEVs derived from dissected MC38 tumor tissues and prove that MLC-V formed based on the tumor tissues TEVs could still inhibit the tumor growth, indicating that MLC-V is a promising strategy as the next generation of personalized therapeutic vaccine. Importantly, the time required for MLC-V manufacturing was about 2 weeks, which is much shorter than 6-18 weeks needed by traditional personalized vaccine procedure. Thus, the application of MLC-V may save both time and cost.

Although anti-PD-1 therapy has shown promising clinical outcomes in colon carcinoma, there is still a hurdle because only a limited number of patients respond^[52,53]^. Importantly, checkpoint inhibition therapy is dependent on the patient’s endogenous tumor-specific CTL responses, and sometimes this is used in combination with other conventional therapeutics to attain optimal efficacy^[54]^. Thus, we investigated the effect of combining anti-PD-1 therapy and found that in MC38 model MLC-V synergize with PD-1 inhibition to elicit much better therapeutic efficacy than each monotherapy. Outstandingly three out of seven mice in combination treatment group were tumor-free, whereas none tumor-free mice were observed in either monotherapy group, suggesting that anti-PD-1 therapy is a useful strategy for augmenting the anti-tumor effect of MLC-V.

Collectively, we demonstrate MLC-V is a time-saving, rapid and low-cost personalized neoantigen-based vaccine manufacturing platform technology. MLC-V can generate desired immune responses in multiple tumor models, and synergize with PD-1 mAb in MC38 model. Thus, MLC-V can be seen as a promising strategy to promote the translation of TEV-based personalized therapeutic vaccine in clinic. However, the detailed mechanism of how PEG-PE interacts with TEVs requires further investigations.

## Author Contributions

W.Z. and W.L. designed the experiments and supervised the study. W.Z., H.T. and Z.W. performed the experiments. H.T. analyzed data. S.L. established the cell line. W.Z. and H.T. composed and edited the manuscript. L.J. coordinated the study and edited the manuscript.

